# Intrinsic structural disorder on proteins is involved in the interactome evolution

**DOI:** 10.1101/2024.02.05.578866

**Authors:** Diego M Bustos

## Abstract

New mathematical tools help to understanding cell functions, adaptability and evolvability to discover hidden variables to predict phenotypes that could be tested in the future in wet labs. Different models have been successfully used to discover properties of the protein-protein interaction network or interactome. We found that in the hyperbolic Popularity-Similarity model cellular proteins with highest contents of structural intrinsic disorder cluster together in many different eukaryotic interactomes and not the prokaryotic *E. coli*, where proteins with high levels of intrinsic disorder are very low. We also found that the normalized theta variable from the Popularity-Similarity model for a protein family correlate to the seniority of the organisms in analysis.

## Introduction

The cell interactome is the sum of all protein connections and the analysis of its structure and evolution is called “comparative interactomics” [1]. The interactome coordinately controls the function of hundreds of proteins [2], and thus influence the evolution of its components. The first high-throughput human protein-protein interaction (PPI) network appeared immediatly after the publication of the human genome, more than two decades ago [3]. Since then, new network-based approaches to PPI analysis have provided novel insights into disease-disease [4] and drug-disease [5] relationships within the human interactome [6]. In addition to this logic, the eukaryotic organism for which we currently have the most complete set of interaction data is the common yeast, *Saccharomyces cerevisiae* [7]. It shares a common ancestor with *Drosophila melanogaster, Caenorhabditis elegans, Mus musculus*, and *Homo sapiens* more than 900 million years ago, probably a distance too large for the retention of similarities in their interaction networks.

PPIs can be described as a set of relationships, encoded in the form of a network in which proteins are represented by nodes and their relationships are represented by connections between these nodes [8]. These networks are driven by variables that are not directly observable or quantifiable and are commonly referred to as hidden or latent variables. The way our data are represented has a fundamental impact on our ability to subsequently extract information from it. To uncover this information, network embedding has proven to be an efficient way to translate graph representations into continuous values [8]. In the Popularity-Similarity model [9], the N nodes (proteins) of a network lie in a hyperbolic space. The hyperbolic geometry is based on parameters different from Euclidean geometry. In this model, the most important variables are the polar coordinates (r_*i*_ and Θ_*i*_). The radial coordinate r_*i*_ represents the popularity or seniority status of node *i* in the system. Nodes that join the system first have more time to accumulate connections (in this case interactions) and are near the center of the circle, while younger nodes are at the edge of the circle and have fewer partners. The angular coordinate Θ_*i*_ allows determining how similar a node *i* is to others. Finally, the hyperbolic distance between nodes, 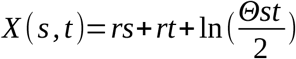 optimizes the process described above, where a new node tries to connect to the most popular node, but also to those most similar to it. Previously, Alanis-Lobato and coworkers embedded the human protein-protein interaction network in a hyperbolic space and found that Θ captures the age and function of the protein [9,10]. In the present work, we link different node properties to the variables (r_*i*_, Θ_*i*_) obtained after embedding PPI from 9 different organisms into the popularity-similarity model. We found a correlation not only with the age, function, and cellular localization of the protein, but also with the degree of structural intrinsic disorder of each protein, the property that it is passed through orthologs. Structural intrinsic disorder is a key property in protein-protein interactions [11,12] because it can allow disordered regions or proteins to structurally accommodate multiple interaction partners [13]. It is defined as the absence of a well-defined three-dimensional structure in proteins. We have observed that r, Θ, and intrinsic disorder correlate between protein families in different organisms. This suggests that evolutionary change in disorder content indicates that disorder evolves to alter function by improving protein stability, regulation, and interactions.

## Methods

### Computer Programming and Statistics

Scripts for data analyses were programmed in Perl and bash. All statistical analyses to evaluate the significance (Wilcoxon rank-sum, Kruskal-Wallis, and Fisher’s exact test) were carried out using the R statistical analysis package. For the analysis of distributions, we used the Wilcoxon rank-sum (paired or unpaired) and Kruskal-Wallis as appropriate.

### Protein interaction network construction

Networks were obtained from the following seven databases BioGRID 4.3.196, DIP, HPRD Release 9, InnateDB 5.4, IntAct 4.2.16, MatrixDB and MINT. All databases were downloaded in 2021, except for the human:yeast inter-interactome which was obtained from [14]. Only physical interactions with the highest percentage of edges supported by more than one experiment were considered. The integrated PINs comprised 29,129 edges and 25,260 nodes for *A. thaliana*; 14,232 edges and 15,972 nodes for *C. elegans*; 44,019 edges and 10,275 nodes for *D. melanogaster*; 26,094 edges and 11,793 nodes for *E. coli*; 249,102 edges and 19,250 nodes for *H. sapiens*; 4,487 edges and nodes for the inter-human:yeast interactome; 34,817 edges and 16,297 nodes for *M. musculus*; 3,698 edges and 15,468 nodes for *R. norvegicus* and 104,110 edges and 6,318 nodes for *S. cerevisiae*. Orthologs were downloaded from Ensembl release 103.

### Intrinsic Disorder Predictions

Intrinsic disorder predictions were done by using the software Espritz [15] (http://protein.bio.unipd.it/espritz/) as previously done in [16,17]. Briefly, the sequence of each protein in the corresponding proteome was downloaded in fasta format from Uniprot (www.uniprot.org). Then Espritz was run on the sequences, resulting files were post-processed using bash scripting.

### Mapping the protein interactomes to hyperbolic space

We embedded the PINs to the hyperbolic space using the R’s plugin LaBNE+HM [10], an approach that combines manifold learning and maximum likelihood estimation to uncover the hidden geometry of complex networks. Once the values of r and Θ were obtained from the package LaBNE+HM those were converted to z score following the formula:

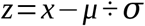

### Gene Ontology enrichment analyses

Gene Ontology enrichment analyses were carried out with the AmigGO webpage [18], and the R package topGO. Only Gene Ontology terms enriched at the 0.05 significance level after Benjamini-Hochberg correction was considered.

### Orthologues

Protein orthologues were detected by using OrthoDB [19] and uniprot [20]. We also identify sequence identity by alignment using Clustal Omega and structural alignment of their PDB stucture when available.

## Results

To investigate hidden biological variables on protein-protein interactome (PPI), we curated the PPI from 9 different interactomes corresponding to the model organisms *Arabidopsis thaliana* (Aa), *Caenorhabditis elegans* (Ce), *Drosophila melanogaster* (Dm), *Escherichia coli* (Ec), *Homo sapiens* (Hs), the interspecies human: yeast (HY) [14,15], *Mus musculus* (Mm), *Rattus norgergicus* (Rn), and *Saccaromise cerevisiae* (Sc).

PPIs are usually considered as scale-free networks [21]. This type of networks is characterized by the presence of hubs. In the interactomes analyzed here, we found strong deviations from strict scale-free behavior, especially in yeast and human PPIs, which are the best studied interactomes. However, in the less studied organisms such as rat, fruit fly, or Arabidopsis, the p(*k*)-versus-*k* diagrams show less deviation from a scale-free network (Fig. 1A). Because the topology of PPI is independent of interaction detection technology [7], it is possible that the observed differences are due to interactome coverage.

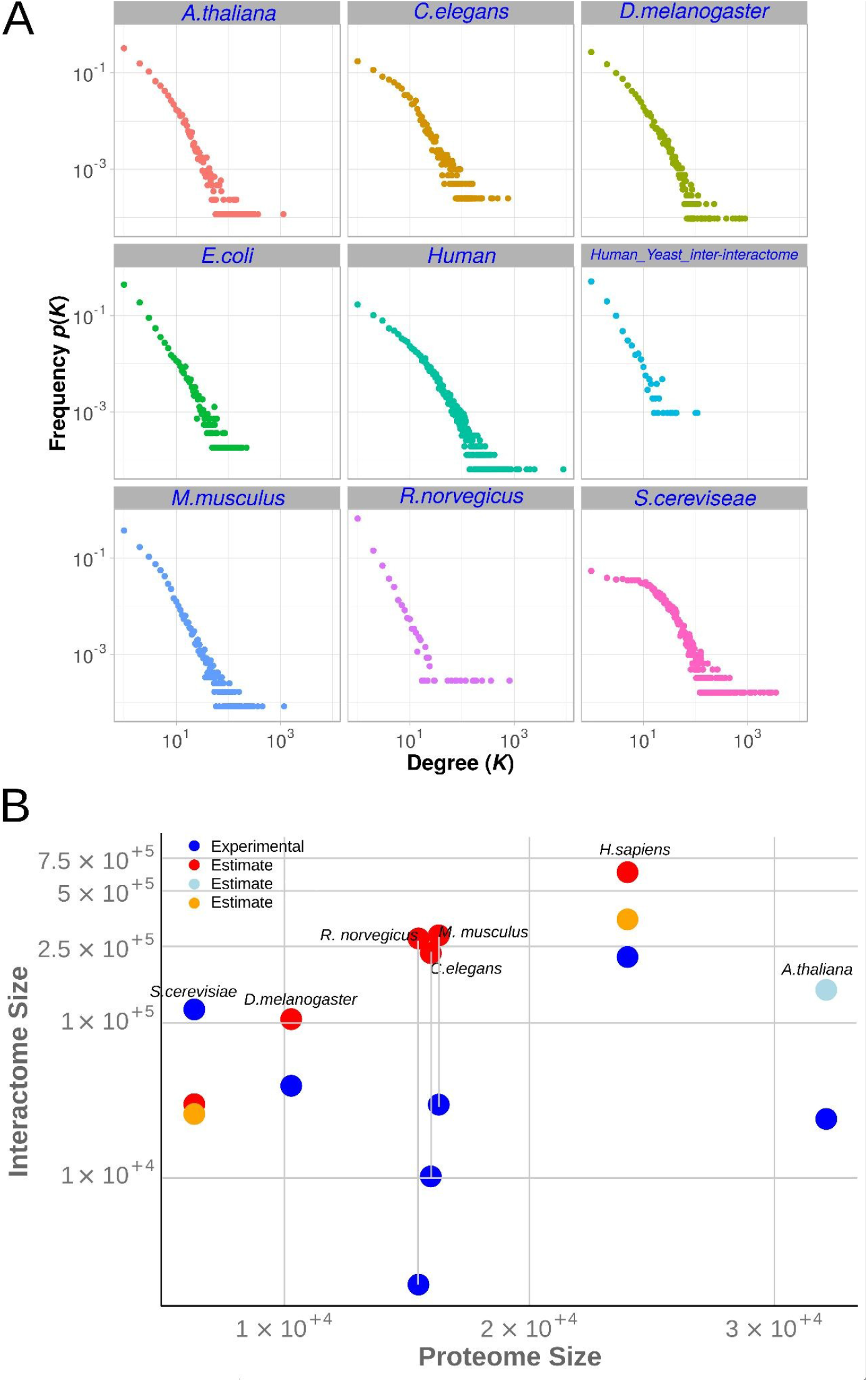

Figure 1B shows the interactome and proteome size for the 9 species. We found that for some proteomes (especially the less studied organisms), the predicted size of the interactome [22] is significantly larger than the available experimental information. In parallel, the relationship between p(*k*) and *k* behaves more like a scale-free network (Fig. 1A) following a power law when the sample size is similar or even larger than the predicted size of the interactome (e.g., in the case of yeast).

The yeast interactome is the only one analyzed here in which the PPI is larger than that predicted on the basis of proteome size [7,22] from the size of its proteome (Fig 1B). It appears that the deviation from the scale-free value is due to the presence of more proteins (nodes) with 10 or fewer partners than expected by theory. To find variables describing the properties of the nodes, we mapped the 9 PPIs to the Popularity-Similarity [9] model proposed by Papadopoulos and Coll. The hyperbolic variable Θ is the most important variable in this model. We mapped many physicochemical properties of proteins and found that proteins with a higher degree of structural intrinsic disorder tended to have lower Θ values than µ (µ is the value of the Z score equal to 0), whereas the more ordered proteins had higher Θ values, as shown in Figure 2.

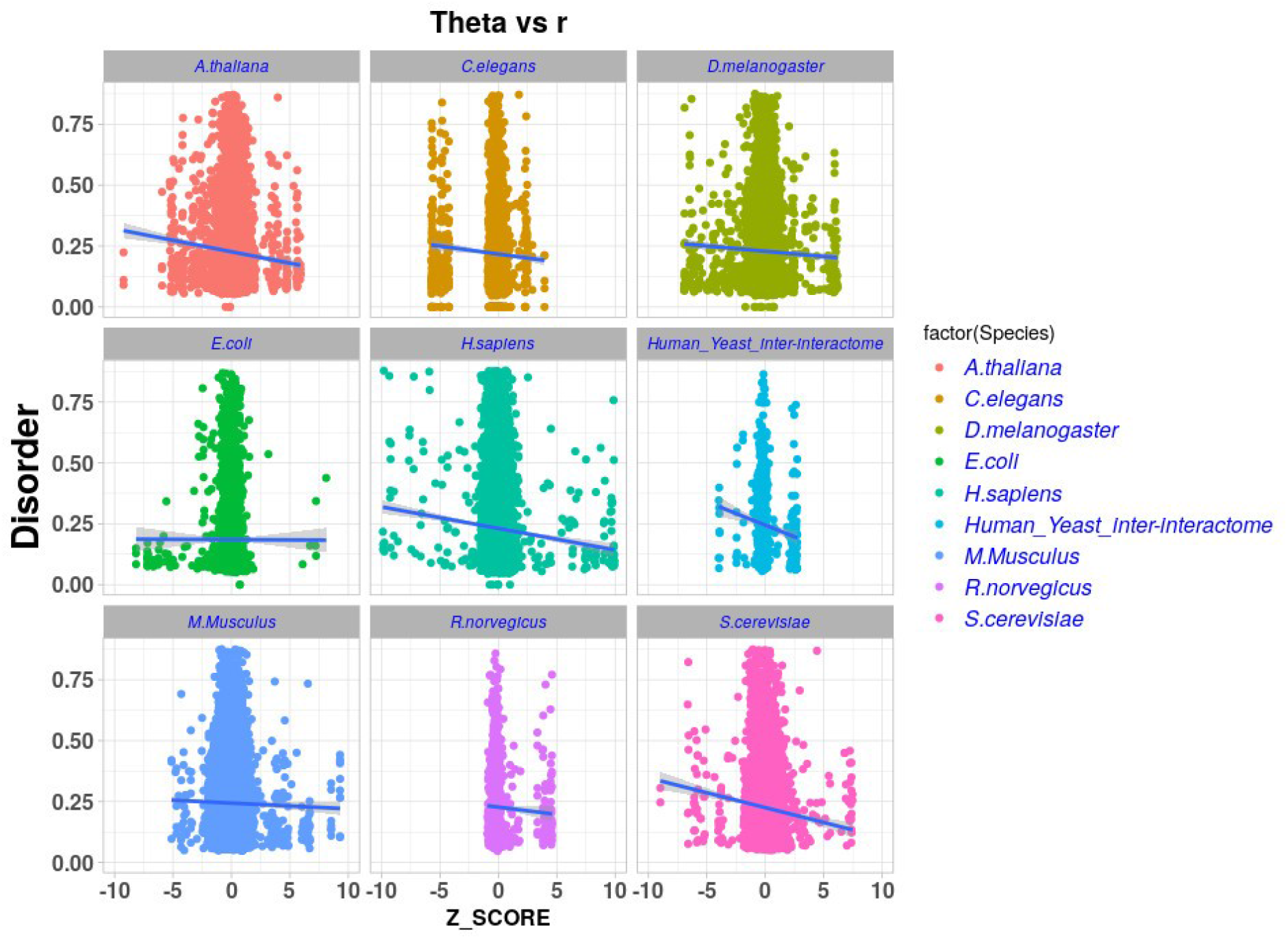

We observed this pattern in all the nine interactomes examined here. Figure 3 shows that the normalized values of Θ statistically capture the sub-cellular localization of proteins (spatial organization of inside the cell) based on protein information from the database Uniprot. However, no enrichment of intrinsic disorder was observed in proteins that share a sub-cellular localization.

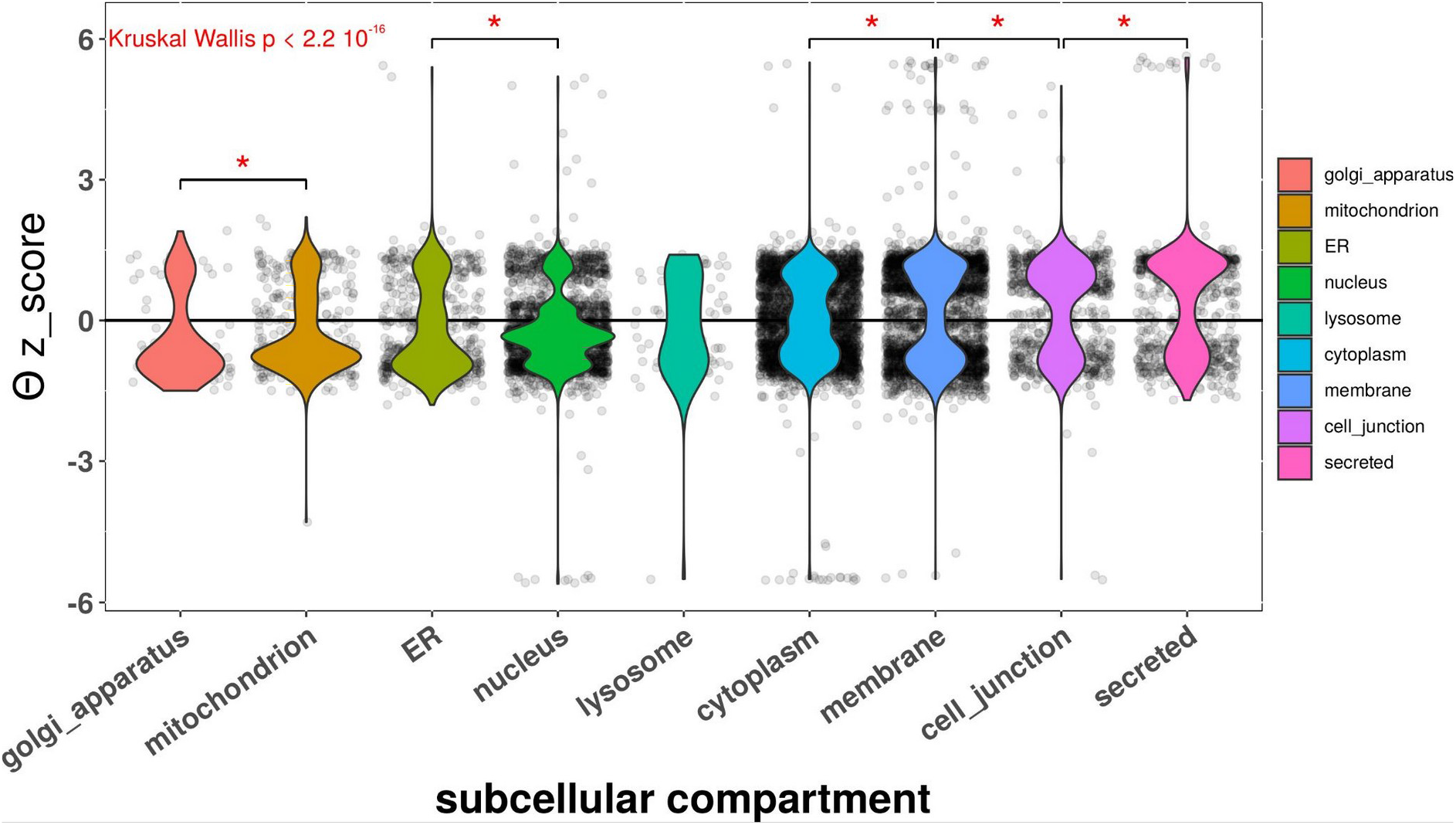

Finally, we analyzed if there is a correlation between the normalized variable Θ and the intrinsic disorder content in a family of orthologues proteins (homologues from different species) from 3 different organisms. We choose human, mouse, and yeast because their relationship and the coverage of their interactomes. We first select protein families from the human interactome with high Θ values (supplementary figure 1), and then we collected their orthologues as described in the method section. We observe that proteins from human have higher normalized Θ values than those from mouse, that at the same time are higher than those for the less organized organism the yeast (Fig 4A). The analyses of the structural disorder of the same proteins show that this characteristic decrease from human to mouse to yeast being statistical significant when human is compared to yeast (Fig 4B). To exemplify this, the figure 4C shows the relation between two tumor suppressors orthologues (P25604_Human and Q99816_Yeast). These proteins are components of the ESCRT-I complex, that is a regulator of vesicular trafficking process. They bind to ubiquitinated cargo proteins and is required for the sorting of endocytic ubiquitinated cargos into multivesicular bodies, and they are required for completion of cytokinesis. In the yeast orthologue the percentage of the protein with high levels of intrinsic disorder in less than 20% and the 80% is a well ordered and structured protein as show in the figure 4C, were the crystal structure of the N and C-term of the protein is shown (PDB ID: 1UZX position 1-161; 2CAZ position 305-385). However, in the human orthologue the 40% of the protein is disordered and only the N-term of the protein could be crystallized (PDB ID: 1S1Q position 1-145).

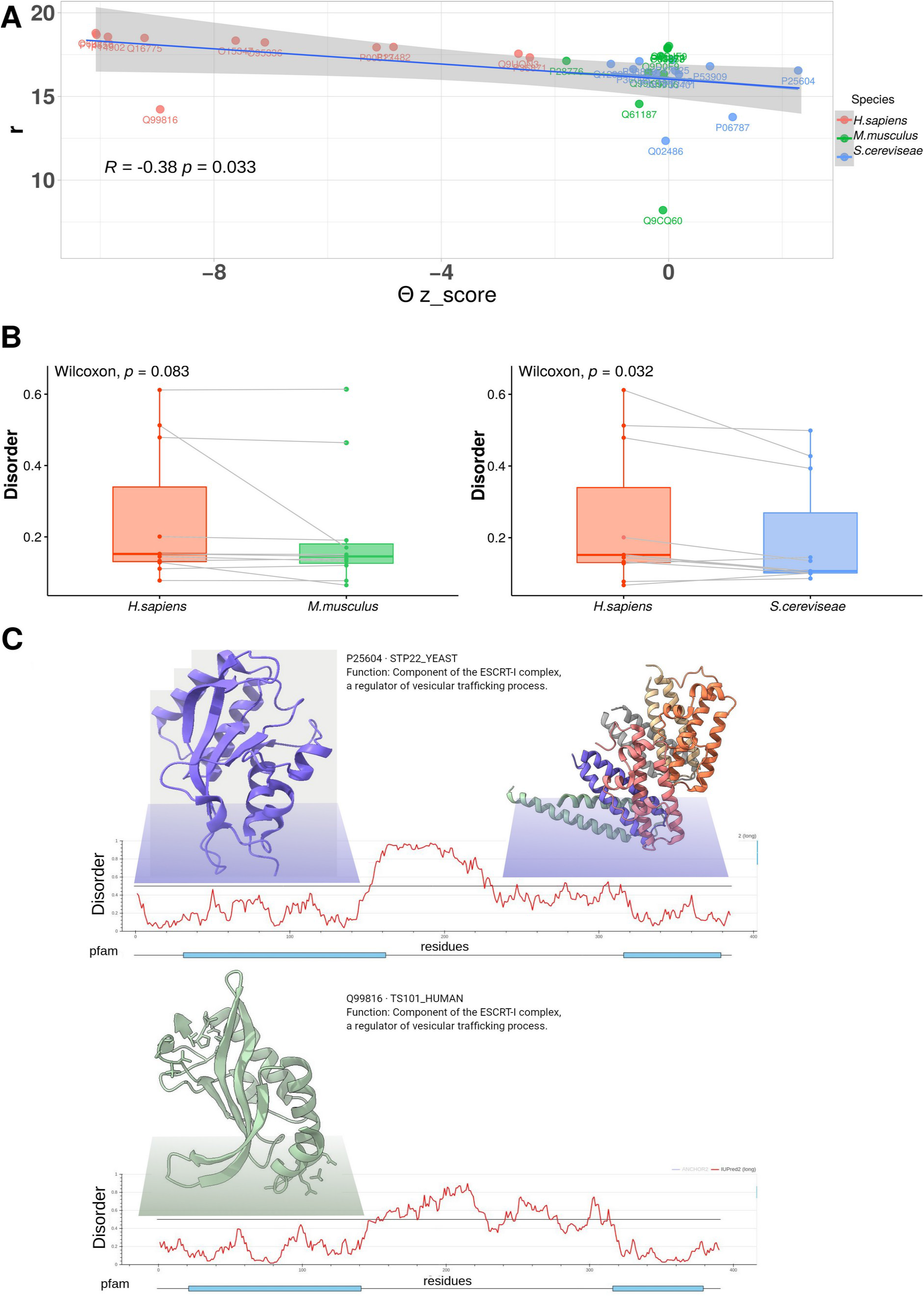

## Discussion

In this study, we try to find hidden variables with correlation to the principal variable in the Popularity-Similarity model of the complex network evolution [9]. After curated large-scales integrated PIN from different model organisms, with a relatively comprehensive coverage, we embedded their in a Popularity-Similarity model to obtain its principal variables. Proteins evolve at different rates along their sequences. In the case of enzymes, rates decreases with the increase of packing, and increases with solvent accessibility [23]. There is a negative correlation between protein sequence evolution and its expression level. This negative correlation triggered the hypothesis of misfolding-driven protein evolution.

that explains the universal dependency between evolution and expression under the assumption that protein misfolding is the principal source of the cost incurred by mutations and errors of translation. This assumption was used to incorporate evolutionary dynamics into an off-lattice model of protein folding. The resulting model of protein evolution reproduced, with considerable accuracy, the universal distribution of protein evolutionary rates, as well as the dependency between evolutionary rate and protein expression. These findings suggest that both universals of evolutionary genomics could be direct consequences of the fundamental physics of protein folding [24].

## Acknowledgments

The authors thank all members of the ISC group for helpful and constructive discussions. This work was supported with funds from CONICET PIP 0118, PUE 0025 and ANPCyT by PICT’17 1984 (Argentina). The authors have no conflict of interest to declare.

